# Latent functional diversity may accelerate microbial community responses to temperature fluctuations

**DOI:** 10.1101/2021.04.14.439774

**Authors:** Thomas P. Smith, Shorok Mombrikotb, Emma Ransome, Dimitrios-Georgios Kontopoulos, Samraat Pawar, Thomas Bell

## Abstract

How complex microbial communities respond to climatic fluctuations remains an open question. Due to their relatively short generation times and high functional diversity, microbial populations harbor great potential to respond as a community through a combination of strain-level phenotypic plasticity, adaptation, and species sorting. However, the relative importance of these mechanisms remains unclear. We conducted a laboratory experiment to investigate the degree to which bacterial communities can respond to changes in environmental temperature through a combination of phenotypic plasticity and species sorting alone. We grew replicate soil communities from a single location at six temperatures between 4°C and 50°C. We found that phylogenetically- and functionally-distinct communities emerge at each of these temperatures, with *K*-strategist taxa favoured under cooler conditions, and *r*-strategist taxa under warmer conditions. We show that this dynamic emergence of distinct communities across a wide range of temperatures (in essence, community-level adaptation), is driven by the resuscitation of latent functional diversity: the parent community harbors multiple strains pre-adapted to different temperatures that are able to “switch on” at their preferred temperature without immigration or adaptation. Our findings suggest that microbial community function in nature is likely to respond rapidly to climatic temperature fluctuations through shifts in species composition by resuscitation of latent functional diversity.

## Introduction

Microbes are drivers of key ecosystem processes. They are tightly linked to the wider ecosystem as pathogens, mutualists and food sources for higher trophic levels, and also play a central role in ecosystem-level nutrient cycling, and therefore, ultimately in global biogeochemical cycles. Temperature has a pervasive influence on microbial communities because of its direct impact on microbial physiology and fitness [1, 2, 3]. There is therefore great interest in understanding how temperature fluctuations impact microbial community dynamics, and how those impacts affect the wider ecosystem [4].

Temperature varies at practically all biologically-relevant timescales, from seconds (e.g., sun/shade), through daily and seasonal fluctuations, to longer-term changes including anthropogenic climate warming and fluctuations over geological timescales. Whole microbial communities can respond to temperature changes over time and space through phenotypic (especially, physiological) plasticity (henceforth, “acclimation”), as well as genetic adaptation in their component populations [5, 6, 7, 8]. Microbial thermal acclimation can occur relatively rapidly (timescales of minutes to days) through processes such as activation and up- or down-regulation of particular genes, and alteration of fatty acids used in building cell walls [9]. Adaptation is a necessarily slower process (timescales of weeks or longer), occurring either through selection on standing genetic variation in the population or that arising through recombination and mutation [5, 10, 11].

In addition, a third key mechanism through which microbial communities can respond to changing temperatures is species sorting [12, 13]: changes in community composition through species-level selection where taxa maladapted to a new temperature are replaced by those that are pre-adapted to it. This can happen either relatively rapidly through the resuscitation or suppression of taxa that are already present [14, 15], or more slowly through immigration-extinction dynamics driven by dispersal from the regional species pool [16, 13]. Resuscitation may be an important mechanism driving species sorting in microbial communities in particular because many microbial taxa have the capacity to form environment-resistant spores when conditions are unfavourable, and then rapidly activate metabolic pathways and resuscitate in favourable conditions. This effectively widens their thermal niche to allow persistence in the face of temperature change [14, 15]. In order for rapid resuscitation of dormant taxa to allow species sorting to drive community-level adaptation, there must be a wide source-pool of species to select from. Indeed, sequencing studies have revealed the presence of thousands of distinct microbial taxa in small environmental samples, most occurring at low abundance [17, 18, 19]. There is also strong evidence that bacteria are often found well outside of their thermal niche. For example, thermophilic taxa are often found in cold ocean beds and cool soils [20, 21, 22]. Thus, a significant reservoir of latent microbial functional diversity may be commonly present for species sorting to act upon [14, 15].

Understanding the relative importance of acclimation, adaptation and species sorting in the assembly and turnover (succession) of microbial communities is key to determining the rate at which they can respond to different regimes of temperature fluctuations. For example, a combination of acclimation and species sorting through resuscitation would enable rapid responses to sudden temperature changes, relative to adaptation. A number of past studies have investigated responses of microbial community composition and functioning to temperature changes, showing that composition can respond rapidly to warming [23, 24], often correlated with responses of ecosystem functioning [25, 26, 27]. However, a mechanistic basis of these community-level responses remains elusive, both in terms of how individual taxa respond to changing temperatures in a community context, and the relative importance of acclimation, adaptation and species sorting. The community context of the responses of individual microbial populations is important because interactions between strains can constrain or accelerate acclimation as well as adaptive evolution [28]. Also, while the importance of species sorting in microbial communities *per se* has been studied [29, 16, 30], work on this issue in the context of environmental temperature is practically non-existent.

A further consideration is whether differing temperature conditions, such as the frequency and magnitude of temperature fluctuations, may influence the life history strategies of the taxa in the community [31, 32], which will in turn alter the relative importance of sorting, acclimation and adaption. In order to identify the life history strategies of bacteria, we must quantify their phenotypic traits, such as growth rates and yield [33]. Quantifying these traits can allow us to identify growth specialists (*r*-strategists) and carrying-capacity specialists (*K*-strategists) [34], and thus test whether these strategies are deferentially favoured in different thermal environments. By identifying life history strategies, we can consider the ecosystem implications of any adaptation-, acclimation-, or sorting-driven changes in microbial communities [33].

Here we investigate whether species sorting and latent functional diversity alone can influence the response of soil bacterial communities to changes in environmental temperature. To this end, we subject replicate communities, shielded from immigration, to a wide range of temperatures in the laboratory. In order to understand the mechanistic basis of observed community-level changes, we analyse the phylogenetic structure and functional traits of the resulting component taxa.

## Materials and Methods

We performed a species sorting experiment to investigate how microbial communities respond to shifts in temperature (Fig. 1). After each community incubation at a given temperature, we estimated the thermal optimum (*T*_opt_) for every isolated strain by measuring the thermal performance curve (TPC) of its maximal growth rate across several temperatures (Fig. 1D). This allowed us to determine how strain-level thermal preferences and niche widths vary with community growth (isolation) temperature, and the presence of taxa pre-adapted to the new temperature. We also performed a phylogenetic analysis of the overall assemblage to identify whether deep evolutionary differences predict which taxa (and their associated traits) are favored by sorting at different temperatures. To quantify strain-level functional traits, we measured their available cellular metabolic energy (ATP), respiration rates, and biomass yield at population steady state (carrying capacity), which allowed us to identify *r*- vs *K*-strategists as well as trade-offs between different strategies.

**Figure 1:**
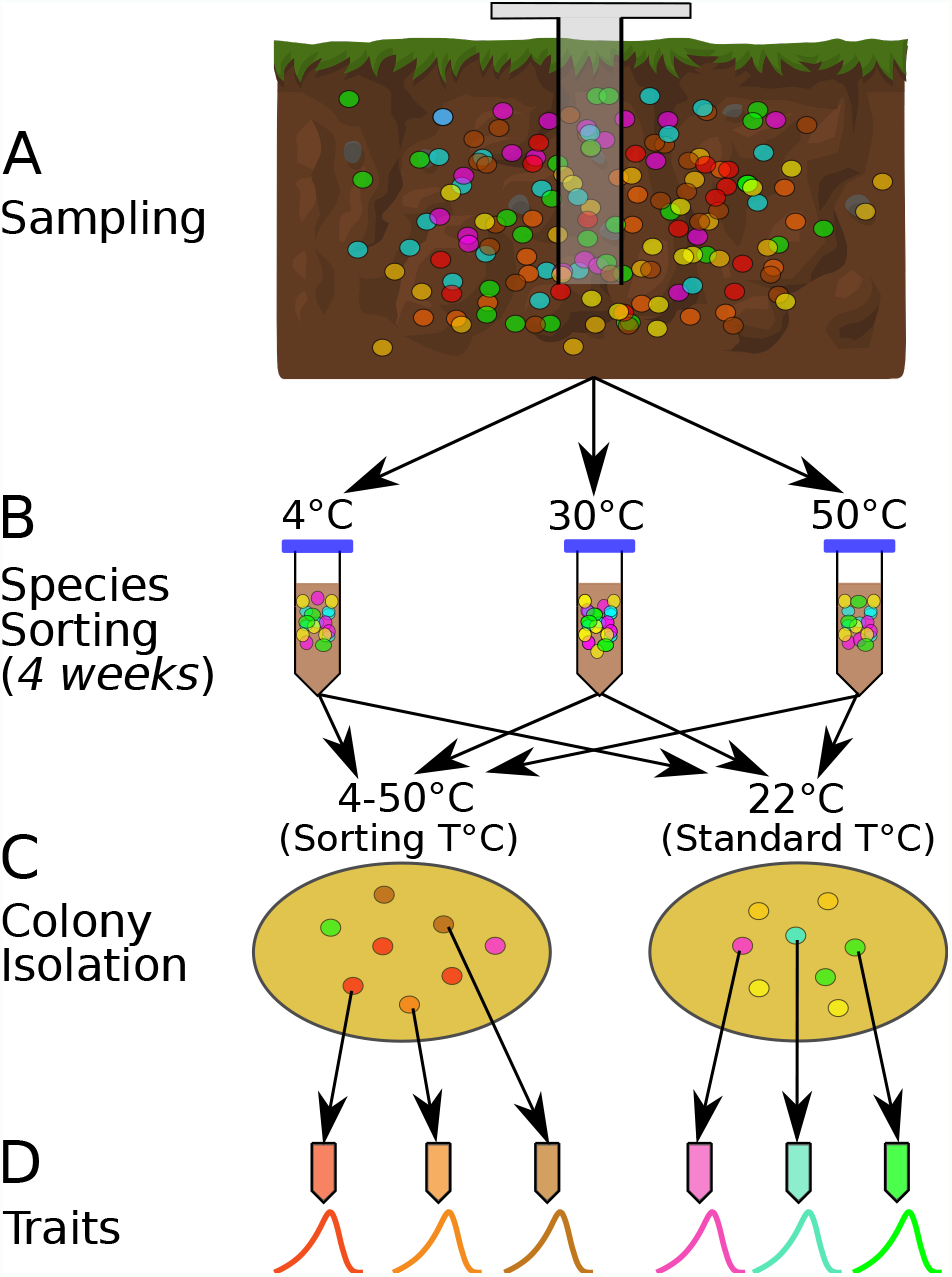
The species sorting experiment. **A:** Different bacterial taxa (coloured circles) sampled from the soil community. **B:** Samples maintained at 4, 10, 21, 30, 40 and 50°C (only three temperatures shown for illustration), allowing species sorting for 4 weeks. **C:** Soil washes from each core plated out onto agar and grown at both the sorting temperature and 22°C (Standard temperature) to allow further species sorting and to facilitate isolation (next step). **D:** The 6 most abundant (morphologically different) colonies from each plate were picked, streaked and isolated, and their physiological and life history traits measured. The curves represent each strain’s unique unimodal response of growth rate to temperature.

### Species sorting experiment

Soil cores were taken from a single site in Nash’s Field (Silwood Park, Berkshire, UK, the site of a long-term field experiment [35]) in June 2016 (Fig. 1A). Six cores were taken from the top 10cm of soil, using a 1.5cm diameter sterile corer. Ambient soil temperature at time of sampling was 19.4°C. The cores were maintained at different temperatures in the laboratory (4°C, 10°C, 21°C, 30°C, 40°C, 50°C) for four weeks to allow species sorting to occur at those temperatures (Fig. 1B). The soil was re-hydrated periodically with sterile, deionised water during incubation. During this period, in each microcosm (incubated soil core), we expected some taxa would go extinct if the temperature was outside their thermal niche, and that survivors would acclimate to the new local thermal conditions. We also expected that the 4-week incubation period would be sufficient time for changes to species interactions due to changes in abundance or traits, and therefore that interaction-driven sorting would occur in addition to the immediate extinctions and acclimation. We then isolated bacterial strains by washing the soil with PBS, plating the soil wash on to R2 agar and incubating the plates at both their 4-week incubation temperature treatments (“sorting temperature”) and at 22°C (“standard temperature”).

The sorting temperature allowed us to determine whether strains in each community tended to have thermal optima matching experimental temperatures, while the standard temperature allowed us to determine whether a 4-week incubation resulted in a loss of taxa that were poorly adapted to 22°C. Appearance of strains with thermal optima matching the standard temperature would indicate incomplete species sorting because the 4-week treatment at temperatures higher or lower than 22°C had not eliminated (or at least suppressed) them.

The plates were incubated until bacterial colonies formed, of which we isolated a single colony from each of the six most abundant morphologically distinct colony types on each plate (Fig. 1C). Additional species sorting likely occurred during this plating-based isolation because strains with the highest growth rates at each temperature would be the first to form visible colonies and be selected. The time-frame for colony appearance on the agar plates differed between temperature treatments, ranging from ∼ 10 days at 4°C to ∼ 1.5 days at 50°C). Morphologically-distinct colonies were isolated from each of the 6 sorting temperatureand 6 standard-temperature plates on R2 agar by streak-plating, before being frozen as glycerol stocks (Fig. 1) which were later revived for trait measurements (see below). In total, 74 strains were isolated in this way.

#### Taxonomic identification

16S rDNA sequences were used to identify the isolates. Raw sequences were first trimmed using Geneious 10.2.2 (https://www.geneious.com), BLAST searches were then used to assign taxonomy to each trimmed sequence. Where all BLAST hits above 97% sequence similarity were to the same species, that species was assigned to the isolate. In cases where no species hits were above 97% sequence similarity, or there were multiple different species hits above 97% similarity, taxonomy was assigned only to genus level.

### Quantifying physiological and life-history traits

#### Growth, respiration, and ATP content

We measured growth rate and respiration rate simultaneously across a range of temperatures for each isolate to construct its acute thermal performance curves (TPCs) for these two traits. We henceforth denote the maximum growth rate across the temperature range by *μ*_max_, and the temperature at which this growth rate maximum occurs, as *T*_opt_ (optimal growth temperature or thermal optimum). ATP content of the entire cell culture was also measured at the start and end of the growth assay. Strains were revived from glycerol stocks into fresh LB broth and incubated to carrying capacity at the temperature of the subsequent experiment. This growth to carrying capacity was an acclimation period, which typically took between 72 hrs (warmest temperatures) to 500 hrs (coldest temperature). Biomass abundance was determined by OD_600_—optical density measurements at 600 nm wavelength. Prior to each growth-respiration assay, the strains were diluted 1:100 in LB, pushing them into a short lag phase before exponential growth started again (also tracked by OD_600_ measurements). The exponentially-growing cultures were subsequently centrifuged at 8,000rpm for 5 minutes to pellet the cells, which were then re-suspended in fresh LB to obtain 400*μ*l culture at a final OD_600_ of ∼ 0.2-0.3. This yielded cells primed for immediate exponential growth without a lag-phase. These cultures were serially diluted in LB (50% dilutions) three times, producing a range of starting densities of growing cells. 100*μ*l sub-cultures of each strain were taken and OD_600_ was tracked in a Synergy HT microplate reader (BioTek Instruments, USA) to ensure that cells were indeed in exponential growth. Initial ATP measurements were made using the BacTiter-Glo assay (see below for details) and cell counts were taken, using a BD Accuri C6 flow cytometer (BD biosciences, USA). Cells were then incubated with a MicroResp™ plate to capture cumulative respiration (see below for details of the MicroResp system) at the experimental temperature and allowed to continue growing for a short period of time (typically 3-4hrs). After growth, the MicroResp plate was read, and final cell count and ATP measurements taken.

We estimated average cell volumes, and calculated the cellular carbon per cell from the flow cytometry cell diameter measurements using the relationship [36]:

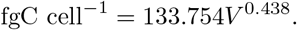

Multiplying this by the cell counts gives an estimate of the carbon biomass of the culture at the starting and ending points.

The difference between the initial biomass and biomass at the end of the experiment gives the total carbon sequestered through growth. Given an initial biomass (*C*_0_) that grows over time (*t*) to reach a final biomass (*C*_tot_), assuming the population is in exponential growth, the mass specific growth rate (*μ*) is given by:

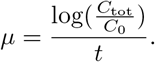

Respiration rates of cultures were measured during growth using the MicroResp™ system [37]. This is a colorimetric assay initially developed to measure CO_2_ production from soil samples, which has since been used to measure respiration of bacterial cultures [38, 39, 40]. We calculate the biomass-specific respiration rate using an equation that accounts for changes in biomass of the growing cultures over time [41]:

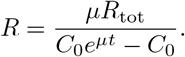

Here, *R*_tot_ is the total mass of carbon produced according to the MicroResp™ measurements, *C*_0_ is the initial population biomass, *μ* is the previously calculated growth rate, and *t* is the experiment duration. ATP content of the cultures was measured using the Promega BacTiter-Glo reagent, which produces luminescence in the presence of ATP, proportional to the concentration of ATP. 50*μ*l of culture (diluted 1:100) was incubated with 25*μ*l reagent. Luminescence was measured over a 6 minute period to allow the reaction to develop completely, and measurements of luminescence recorded at the 0, 2, 4 and 6 minute timepoints. The highest relative light units (RLU) measurement for each culture was taken and used to calculate the quantity of ATP, using

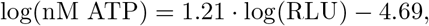

derived from a calibration curve. This was then normalised by the flow cytometry measurements to calculate a value of ATP/biomass.

#### Thermal performance curves

To quantify thermal performance curves of individual isolates, we fitted the Sharpe-Schoolfield model with the temperature of peak performance (*T*_pk_) as an explicit parameter [42, 43] to the experimentally-derived temperature-dependent growth rate and respiration rates of each isolate:

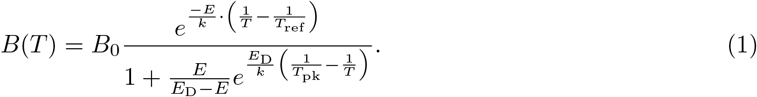

Here, *T* is temperature in Kelvin (K), *B* is a biological rate (in this case either growth rate, *μ*, or respiration rate, *R*), *B*_0_ is a temperature-independent metabolic rate constant approximated at some (low) reference temperature *T*_ref_, *E* is the activation energy in electron volts (eV) (a measure of “thermal sensitivity”), *k* is the Boltzmann constant (8.617 × 10^−5^ eV K^−1^), *T*_pk_ is the the temperature where the rate peaks, and *E*_D_ the deactivation energy, which determines the rate of decline in the biological rate beyond *T*_pk_. We then calculated the peak performance (i.e., *R*_max_ or *μ*_max_) by solving Eq. 1 for *T* = *T*_pk_. This model was fitted to each dataset using a standard non-linear least squares procedure [41].

The *T*_pk_ for growth rate was considered the optimum growth temperature (i.e., *T*_opt_) for each isolate. Then, the operational niche width was calculated as the difference between *T*_opt_ and the temperature below this value where *μ*_max_ (*B*(*T*) in eqn 1) reached 50% of its maximum (i.e, *μ*_max_ at *T*_opt_). This, a measure of an organism’s thermal niche width relevant to typically experienced temperatures [44, 8], was used as a quantification of the degree to which an isolate is a thermal generalist or specialist.

In most cases, *T*_opt_ was derived directly from the Sharpe-Schoolfield flow-cytometry growth rate fits. Four strains of *Streptomyces* were unsuitable for standard flow-cytometry methods due to their formation of mycelial pellets [45]. For these strains, growth rates derived from optical density measurements were used to estimate *T*_opt_ instead. Finally, for six isolates whose growth was recorded at too few temperatures to fit the Sharpe-Schoolfield model, the temperature with the highest directly-measured growth rate was taken as an estimate of *T*_opt_.

#### Trade-offs between traits

To understand the trade-offs and collinearities between different life history and physiological traits, we performed a principal components analysis (PCA), with optimum growth temperature (*T*_opt_), niche width, peak growth rate (*μ*_max_), peak respiration rate (*R*_max_), mean cellular ATP content (log-transformed), and carrying capacity (OD_600_) as input variables (scaled to have mean = 0 and s.d. = 1).

All rate calculations, model fitting and analyses were performed in R.

### Phylogenetic trait mapping

We used the 16S sequences to build a time-calibrated phylogeny in order to investigate the evolution of thermal performance across the isolated bacterial strains. Sequences were aligned in MAFFT (v7.205), a tree was built with RAxML (v8.1.1), and time-calibration was performed using PLL-DPPDiv [46] (see Supplementary Information for full details).

To test whether there was evidence of evolution of *T*_opt_, we calculated Pagel’s *λ* [47], which quantifies the strength of phylogenetic signal — the degree to which shared evolutionary history has driven trait distributions at the tips of a phylogenetic tree. *λ* = 0 implies no phylogenetic signal, i.e. the signal expected if variation in trait values is independent of the phylogeny. *λ* = 1 implies strong phylogenetic signal, i.e. that the trait has evolved gradually along the phylogenetic tree (approximated as Brownian motion). Intermediate values (0 *< λ <* 1) imply deviation from the Brownian motion model, an may be observed for different reasons, such as constrained trait evolution due to stabilizing selection, variation in evolutionary rate over time (e.g., due to episodes of rapid niche adaptation). Pagel’s *λ* requires that the trait be normally distributed. However *T*_opt_ in our dataset has a right-skewed distribution. Therefore, to test phylogenetic heritability we calculated *λ* for log(*T*_opt_).

We mapped the evolution of *T*_opt_ onto our phylogeny using maximum likelihood to estimate the ancestral values at each internal node, assuming a Brownian motion model for trait (an appropriate model, given the obtained *λ* value).

The estimates of phylogenetic signal and the visualisation of trait evolution were performed using tools from the R packages ape and phytools [48, 49]. The p-value for phylogenetic signal was based on a likelihood ratio test.

## Results

### Species sorting

In total, 74 strains of bacteria were isolated; six from each incubation temperature with matching sorting isolation temperature and six from each incubation temperature followed by a standard isolation temperature, with the exception of the 30°C sorting temperature regime, from which we obtained 8 isolates. Of these isolates, 62 could be reliably revived in liquid culture, from which 53 grew across a wide enough temperature range to produce enough data points for fitting the Sharpe-Schoolfield model (Eq. 1). The 62 strains that could be revived were from 14 genera within 3 bacterial phyla (Supplementary Table S1). Isolates were in general well adapted to their sorting temperature (Fig. 2A). A quadratic linear regression model fitted the data well (p *<* 0.0001, shown in Fig. 2A) and was preferred to a straight-line regression model (ANOVA, p *<* 0.0001). The deviation from a simple linear response arises because the *T*_opt_’s of isolates from the three lowest temperatures (4°C, 10°C, and 21°C) are significantly higher than their sorting and isolation temperature (Fig. 2A). In comparison, the *T*_opt_’s of standard temperature isolates were largely independent of the temperatures that their community had been previously grown at (Fig. 2B), indicating that species sorting of the 4-week period had been incomplete, i.e., strains maladapted to those temperature treatments had not been eliminated and were able to be resuscitated.

**Figure 2:**
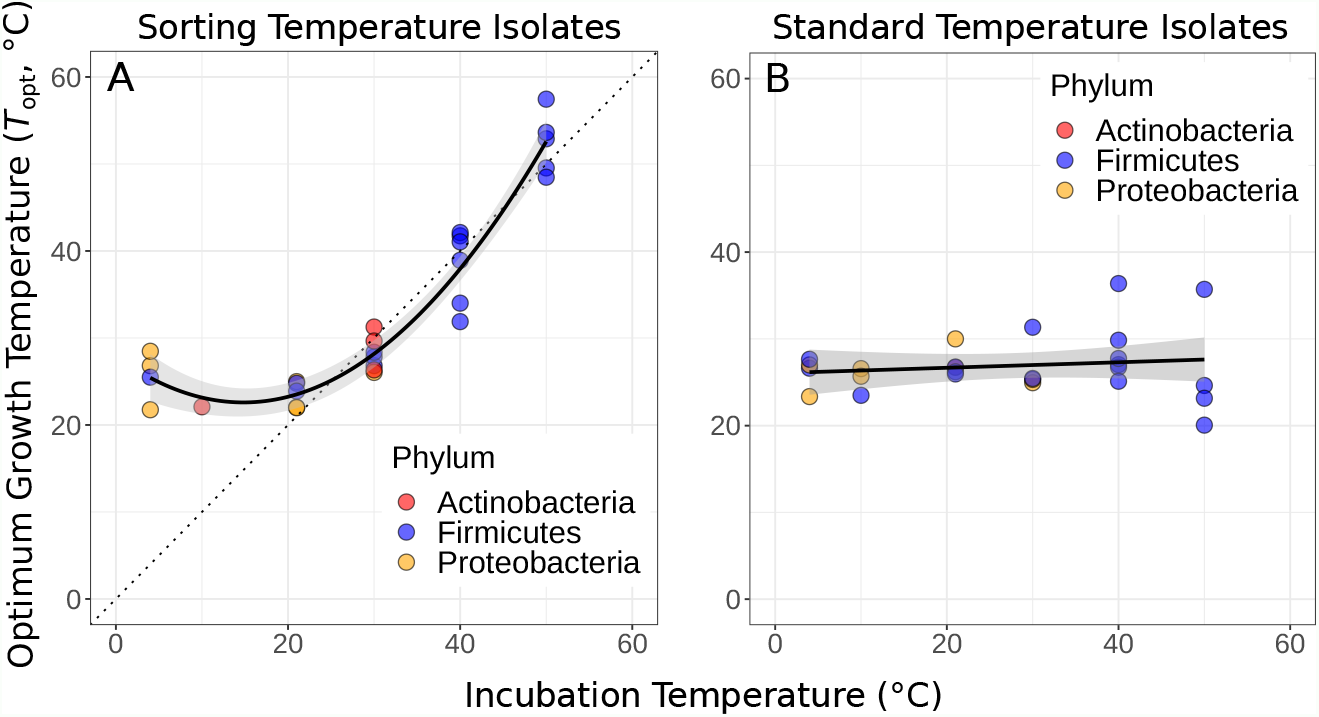
Species sorting of soil bacteria driven by temperature change. **A:** Thermal optima of growth rate closely match sorting temperature for the isolates from those temperatures (black line: quadratic linear regression, p *<* 0.0001, R^2^ = 0.66). Note that the prediction bounds at three lowest temperatures do not include the 1:1 (dashed) line. **B:** No significant association between incubation temperature and thermal optima for standard temperature isolates (simple linear regression, p = 0.74). These results show that species sorting can act upon latent diversity to select for isolates adapted to different temperature conditions (**A**), but that isolates maladapted to the sorting conditions can reemerge (be resuscitated) under the appropriate conditions (**B**).

### Evolution of *T*_opt_

*T*_opt_ displays a strong signal of phylogenetic heritability, closely approximating a Brownian motion model of trait evolution (Pagel’s *λ* = 0.99, p < 0.001), i.e. closely related species have more similar *T*_opt_ than random pairs of species. Qualitatively the same result was obtained using Blomberg’s *K* metric (cf. Supplementary Information Section 4). The estimated ancestral states of *T*_opt_ were mapped onto a phylogeny, where it can be seen that colder- or hotter-adapted strains tend to cluster together (Fig. 3A). The inferred evolution of *T*_opt_ through time indicates that a large amount of the trait space (cool to hot) is explored by *Firmicutes*, while *Actinobacteria* and *Proteobacteria* are constrained to a much narrower range of (relatively cool) optimal growth temperatures (Fig. 3B).

**Figure 3:**
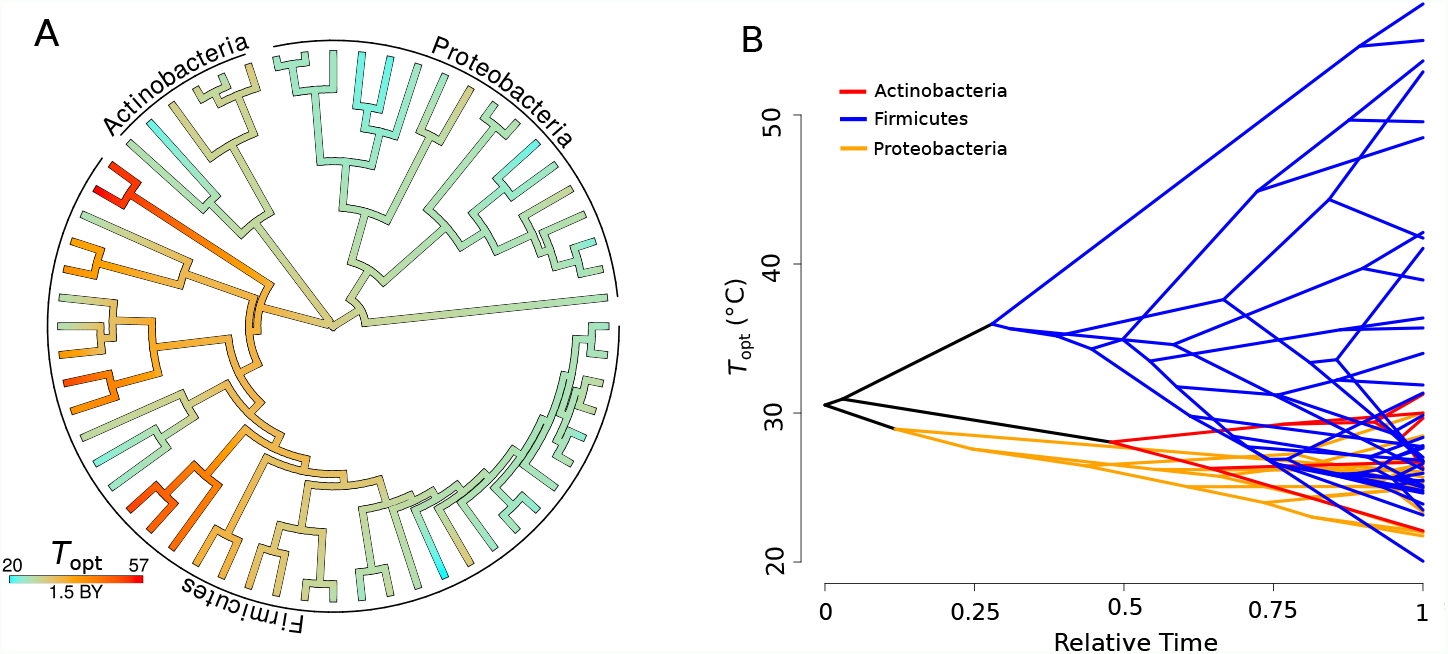
Evolution of *T*_opt_. **A**. Ancestral trait reconstruction of *T*_opt_ visualised on a tree, from lower temperatures in cyan, to higher temperatures in red, with time given in billions of years (BY). All of the higher temperature (40-50°C) isolates belong to the phylum *Firmicutes*. **B**. Projection of the phylogenetic tree into the *T*_opt_ trait space (y-axis), over relative time (x-axis) since divergence from the root. The clades representative of each phylum are coloured on the projection (*Actinobacteria* red, *Firmicutes* blue, *Proteobacteria* yellow).

### Functional traits and life history strategies

We investigated the level of association and trade-offs between different traits in the two major phyla isolated (*Firmicutes* and *Proteobacteria*) using PCA (Fig. 4A). Growth specialists (copiotrophs, *r*-specialists) are expected to grow rapidly but wastefully, and therefore have high ATP content in combination with high growth rates, but low overall yield (carrying capacity). Yield specialists (oligotrophs, *K*-specialists) are expected to grow more slowly but more efficiently, and should therefore display the opposite pattern, i.e. relatively low growth rates and ATP content, but high yield. The first two principal components explained 60.1% of the cumulative variation in the data. *T*_opt_, carrying capacity and respiration rate showed greatest loading on the first principal component (PC1), whilst growth rate and niche width load most strongly on PC2. The *Firmicutes* and *Proteobacteria* phyla are partitioned in this space. The positive loadings onto PC2 of growth rate and ATP content versus the negative loading of carrying capacity suggest an *r* vs *K* growth strategy trade-off; *Proteobacteria* have traits associated with a *K*-selected life history strategy while *Firmicutes* tend to have traits associated with a *r*-selected strategy. Furthermore, thermal niche width loads positively on PC2 along with growth rate and ATP content, implying that thermal generalism is not traded off against growth rate in these taxa — i.e., no thermal generalist-specialist trade-off in growth rates.

**Figure 4:**
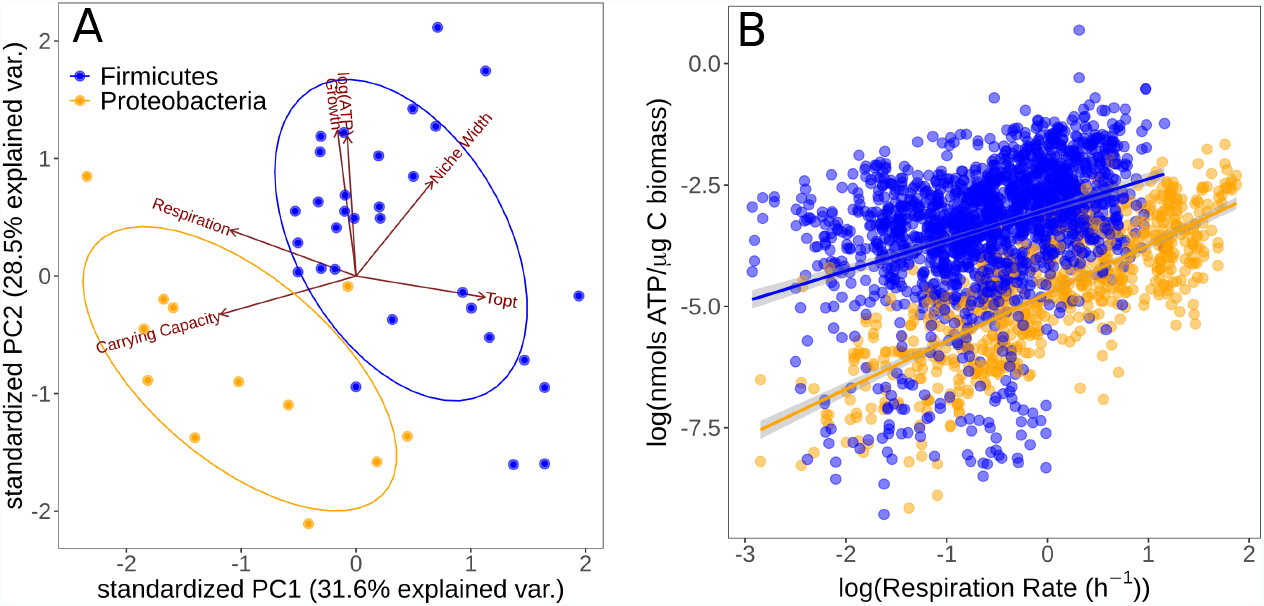
Partitioning of growth strategies between phyla. **A:** PCA on life-history traits, coloured by phylum. Relative to each other, *Firmicutes* (blue) tend to be *r* specialists, *Proteobacteria* (orange) tend to be *K* specialists. **B:** ATP content of cultures is associated with the respiration rate. *Firmicutes* show a sub-linear scaling relationship of ATP with respiration rate (scaling exponent = 0.60 ± 0.07), whilst *Proteobacteria* display an approximately linear scaling relationship (scaling exponent = 0.99 ± 0.06). The same colour scheme is shared by both sub-plots.

To further understand the partitioning of taxa into these life-history strategies, we investigated the differences in accessible cellular energy (ATP) content between these two phyla. We found that across the entire dataset, respiration rate and ATP content display a power-law relationship in both phyla (Fig. 4B). Whilst *Firmicutes* have generally higher ATP levels overall, they display a sub-linear scaling relationship of ATP levels with respiration rate (scaling exponent = 0.60 ± 0.07). In comparison, whilst *Proteobacteria* have less standing ATP content on average, they show an approximately linear scaling relationship between ATP and respiration rate (scaling exponent = 0.99 ± 0.06). This suggests that *Proteobacteria* are deriving ATP from aerobic respiration only, whereas *Firmicutes* may be utilising alternative pathways.

## Discussion

Here, using a novel species sorting experiment, we have studied the extent to which species sorting and acclimation can influence the responses of soil bacterial communities to temperature change. We find that when replicate soil bacterial communities sampled from a temperate environment are subjected to a wide range of temperatures for 4 weeks, in microcosms where immigration is not possible, strains with thermal preferences matching the local conditions emerge consistently. The strong correspondence between strain-level optimal growth temperatures and isolation temperatures (Fig. 2A) indicates that a pool of taxa with disparate thermal physiologies, including those maladapted to the ambient thermal conditions, persisted in the parent community. This result is reinforced by fact that the *T*_opt_’s of standard temperature isolates were largely independent of the temperatures that their community had been previously grown at (Fig. 2B), indicating that strains maladapted to that temperature had not been eliminated and were able to be resuscitated.

While a 4 week period is arguably too short for mutation or recombination driven thermal adaptation in environmental samples (as a significant degree of generational turnover is required [5, 50, 51]), it is worth considering the possibility that some of the community-level emergence of thermally-adapted strains could have been driven by rapid evolution through selection on standing trait variation. Indeed, stochastic mapping of thermal physiological traits on the prokaryotic tree of life has shown that *T*_opt_ evolves relatively rapidly compared to other traits such as niche width or activation energy (thermal sensitivity) [8]. This is consistent with adaptive evolution experiments, which have shown that bacteria as well as archaea can rapidly adapt to new temperatures by shifting their *T*_opt_ [5, 6, 7, 3]. The molecular mechanisms underlying such rapid evolution are still being investigated, but structural changes to enzymes that alter their melting temperatures appears to be a key mechanism when adaptation to relatively high temperatures is called for [52]. While determining whether such mechanisms can be operationalised over the duration of our sorting experiment was beyond the scope of our study, this is still arguably a very short time-frame for significant shifts in thermal optima due to selection on standing variation alone. Furthermore, the communities that remained after 4 weeks of growth at the six temperatures consisted of taxonomically-distinct sets of strains, and the *T*_opt_’s of the overall set of taxa exhibited a significant phylogenetic signature (Fig. 3). This indicates that the observed systematic differences in *T*_opt_ across the temperature-specific communities were driven by species sorting on pre-existing physiological variation across strains, rather than thermal adaptation of single strains. Overall, we therefore conclude that species sorting played a dominant role in determining the response of the parent community to abrupt changes in temperature, in the absence of immigration, and with negligible adaptation.

We also detected systematic turnover in functional traits that likely underpin the change in thermal optima with species sorting. There were differences in the taxa isolated at different temperatures, with more *Proteobacteria* at lower temperatures and more *Firmicutes* at higher temperatures (all *T*_opt_ *>* 35°C were *Firmicutes*). Furthermore, these phyla were partitioned in the *r*-*K* and thermal generalism-specialism trait spaces (Fig. 4). *Proteobacteria* were found to be relatively *K*-selected thermal specialists and *Firmicutes* relatively *r*-selected thermal generalists. These findings are inconsistent with a generalist-specialist trade-off in which increasing thermal niche width is proposed to inevitably incur a metabolic cost, reducing maximal growth rates [53, 54]. As with our findings, recent work on phytoplankton thermal performance traits also failed to detect a generalist-specialist trade-off [43], questioning its universality in microbes. Since the existence of such a trade-off plays a key role in life history theory, there would be value in further experiments to confirm the generality of this finding.

The increased growth rates and lower respiration rates of *Firmicutes* relative to *Proteobacteria* are also consistent with data from meta-analyses of bacterial rates [55, 3] (cf. Supplementary Fig. S1). Additionally, previously reported values for cellular ATP content are generally found to been higher for *Firmicutes* than *Proteobacteria* with more than 10-fold greater intracellular ATP content reported for *Bacillus* versus *Pseudomonas* strains [56], some of the major representatives of *Firmicutes* and *Proteobacteria* in this experiment respectively. This suggests that these phyla tend to allocate resources to growth and respiration in fundamentally different ways. One explanation for these seemingly divergent strategies may be found in *Firmicutes* deriving extra energy though fermentation pathways. There is a mechanistic trade-off between growth rate and yield whereby bacteria may increase their rate of ATP production by supplementing aerobic respiration with fermentation [57]. Fermentation pathways increase the rate of ATP production but result in lower total yield, allowing populations to reach higher growth rates but lower carrying capacity from the same resource input. This is consistent with the apparent *r* vs *K* selection trade-off observed in our results. The differences in scaling relationship between ATP content and respiration rate may provide further evidence of differences in metabolic pathways utilised. Across *Proteobacteria*, ATP content has a scaling exponent of approximately 1 with respiration rate, indicating that theses strains are deriving ATP solely from aerobic respiration (Fig. 4B). The fact that *Firmicutes* have a lower scaling exponent (0.60 ± 0.07) i.e., that they are generating higher levels of ATP than expected at lower rates of respiration, may indicate that they derive ATP from alternative pathways alongside aerobic respiration. These differences in metabolic strategies reflect underlying differences in the efficiency of growth, *i*.*e*. carbon use efficiency (CUE), between these taxa [41]. Moreover, CUE varies systematically with temperature, in a phylogenetically structured manner [58, 41]. Thus, community turnover due to temperature change is likely to have a profound impact on community-level functional traits, such as CUE [59].

In contrast to the strong association between *T*_opt_ and incubation temperature in the sorting temperature isolates, we did not observe any similar relationships in the standard temperature isolates, where mesophiles were consistently recovered regardless of prior incubation conditions. This indicates that species sorting was incomplete (in that maladapted taxa were not driven extinct), implying that bacterial communities can be resilient to temperature change at the community level. Taxa suited to different temperatures are able to “switch on” as conditions become suitable, allowing community-level functional plasticity due to the latent functional diversity present within communities. Although mesophiles were recovered from all incubation temperatures in our standard temperature experiment, there was the same taxonomic bias as seen in the sorting temperature isolates – more *Firmicutes* were recovered from higher temperatures. This is probably a reflection of the propensity of Firmicutes to form endospores and remain dormant until conditions are favourable, upon which they invest resources into rapid growth to gain a competitive advantage over other taxa, consistent with our life-history trait findings of *r*-specialism in *Firmicutes* [14]. In comparison, the *Proteobacteria* in this experiment were generally more suited to oligotrophic environments (e.g. *Collimonas*, [60]), where constituent species are expected to present low growth rate and high carrying capacity (*K* specialists, [61]) as well as increased respiration [62]. This idea is supported by the observation that we isolated the strains from sandy, acidic soil (i.e. oligotrophic) [63] and sequencing studies revealed that *Proteobacteria* are the most abundant phylum [35]. We do not suggest that this adoption of *r* versus *K* strategy is general to all *Firmicutes* and *Proteobacteria*. Indeed meta-analysis reveals little consistency in the phyla associated with copiotrophy or oligotrophy [64]. Nor do we suggest that warming is likely to result in selection for *Firmicutes* over *Proteobacteria* – community temperature responses are not likely to be consistent at coarse phylogenetic levels [1]. However, the results presented here are consistent with phylum-specific traits for the majority of our isolates, when compared to each other.

Patterns of microbial community succession in nature are driven by the differences in growth strategy between taxa that we report here. Studies have revealed taxonomic groups associated with different stages of microbial succession, with patterns broadly consistent across timescales of days [65, 66, 67] and years [68, 69] and even over thousands and tens of thousands of years, as revealed through sequencing of soil sediments [70]. Generally, across these studies, the phyla *Firmicutes* and *Bacteroidetes* are associated with early succession, whilst other phyla such as *Actinobacteria* and *Acidobacteria* are more abundant at later stages of succession. *Proteobacteria* are less consistent at the phylum level, with *Alphaproteobacteria* associated with late succession, *Betaproteobacteria* associated with early succession and *Gammaproteobacteria* variously associated with different stages of succession in different studies. Isolated taxa reveal a strong association between early succession and high growth rates [66] as well as rRNA operon copy numbers, a key determinant of bacterial growth rate [71]. The *K*-selected taxa may therefore be thought of as general constituents of soil, associated with standard low turnover of carbon, whilst the *r*-selected taxa may been seen as more opportunistic from their involvement in early succession. Indeed, signatures of community-level differences in *r*-versus *K*-selection have been observed in microbial communities at different successional stages [72]. Fluctuating temperatures may therefore drive repeated assembly dynamics via sorting on latent microbial diversity, leading to functional community changes through time. However, the frequency and magnitude of temperature fluctuations may also influence the life history strategies of the taxa in the community [31, 32].

In summary, we have found that resuscitation of latent functional diversity driven by phenotypically plastic responses of single taxa to temperature change can allow whole bacterial communities to track dramatic changes in temperature. Community function is expected to be driven by interactions between the most abundant taxa [73] and therefore changes in the abundance of taxa with temperature variation are likely to drive profound changes in overall community functioning (mediated by community-level variation in traits such as CUE). In particular, *r*-vs *K*-selection is likely to vary with temperature change at the community level, from daily to seasonal successional trajectories, driven by species sorting. Furthermore, climate change is expected to lead to increased temperature fluctuations [74], both in magnitude and frequency. This may potentially lead to more frequent species sorting effects over short timescales, further driving changes in community composition through time. Overall, these results show that latent diversity in thermal physiology combined with temperature induced species sorting is likely to facilitate the responses of microbial community structure and functioning to climatic fluctuations.

## Acknowledgments

TPS was supported by a BBSRC DTP scholarship (BB/J014575/1). TB and SP were funded by NERC grants NE/M020843/1 and NE/S000348/1.

