## Supplementary Information for "Latent functional diversity may accelerate microbial community responses to temperature fluctuations"

### Supporting Information for
### Latent functional diversity may accelerate microbial community responses to temperature fluctuations

Thomas P. Smith, Shorok Mombrikotb, Emma Ransome, Dimitrios-Georgios Kontopoulos, Samraat Pawar, Thomas Bell

#### Contents

**1 Details of isolated strains 1**

**2 Phylum differences seen in alternative datasets 3**

**3 Phylogenetic reconstruction 3**

**4 Alternative model of trait evolution 4**

**5 Timetree calibration nodes 4**

#### 1 Details of isolated strains

Full details of all the isolates and their taxonomy as determined through 16S sequencing is shown in Table S1. Whilst there is taxonomic diversity in the isolates, there were also genetically similar isolates (tentatively the same "species" or "ecotypes") which were obtained more than once. In particular, isolates from the *Bacillus cereus* group, which consists of several genetically similar species - *B. anthracis*, *B. cereus*, *B. mycoides*, *B. thuringiensis* and *B. weihenstephanensis* [Logan and Vos, 2015], were commonly found across many temperature treatments. Their close genetic relatedness prevents 16S rDNA sequences from being a good tool for accurate species delimitation [Ash et al., 1991], so taxonomy was assigned based on phenotypic characteristics. *B. mycoides* were differentiated from other *Bacillus cereus* groups strains by rhizoid colony architecture [Logan and Vos, 2015]. Strains displaying cold tolerance (growth at 5°C) were designated as *B. weihenstephanensis* [Lechner et al., 1998, Logan and Vos, 2015]. Finally, two strains with highest BLAST matches to *B. cereus*, *B. mycoides* and *B. weihenstephanensis* but displaying neither cold tolerance, nor rhizoid colonies, were designated *B. cereus*. Due to the high similarity of 16S sequences across this group, these strains do not cluster into monophyletic species groups.

Table S1: **List of strains and ecotypes isolated.**

Strain codes follow XX_YY_ZZ naming convention, where XX is the incubation temperature, YY is the isolation temperature and ZZ is a numeric designator for the specific isolate. RT = room temperature (22°C, termed "standard temperature" in the main text).

| Ecotype | Associated Strains | Phylum | Class | Order | Family |
|---|---|---|---|---|---|
| Anoxybacillus caldiproteolyticus | 50_50_06 | Firmicutes | Bacilli | Bacillales | Bacillaceae |
| Anoxybacillus tepidamans | 50_50_02 | Firmicutes | Bacilli | Bacillales | Bacillaceae |
| Arthrobacter sp. | 10_10_06 | Actinobacteria | Actinobacteria | Micrococcales | Micrococcaceae |
| Bacillus bataviensis | 40_RT_02 | Firmicutes | Bacilli | Bacillales | Bacillaceae |
| Bacillus megaterium | 40_40_01; 40_40_03; 21_RT_02; 40_RT_05 | Firmicutes | Bacilli | Bacillales | Bacillaceae |
| Bacillus mycoides | 21_21_05; 30_30_05; 30_RT_03 | Firmicutes | Bacilli | Bacillales | Bacillaceae |
| Bacillus cereus | 21_21_01; 21_R_06 | Firmicutes | Bacilli | Bacillales | Bacillaceae |
| Bacillus weihenstephanensis | 04_04_05; 30_30_02; 30_30_04; 04_RT_01; 04_RT_03; 10_RT_03; 30_RT_06; 40_RT_06; 50_RT_02; 50_RT_06 | Firmicutes | Bacilli | Bacillales | Bacillaceae |
| Bacillus simplex | 40_RT_01; 50_RT_01 | Firmicutes | Bacilli | Bacillales | Bacillaceae |
| Bacillus sp. bataviensis-like | 50_RT_04 | Firmicutes | Bacilli | Bacillales | Bacillaceae |
| Bacillus sp. niacini-like | 50_RT_03 | Firmicutes | Bacilli | Bacillales | Bacillaceae |
| Bacillus sp. Shackletonii-like | 50_50_05 | Firmicutes | Bacilli | Bacillales | Bacillaceae |
| Brevibacillus thermoruber | 50_50_03; 50_50_04 | Firmicutes | Bacilli | Bacillales | Bacillaceae |
| Cohnella sp. | 40_40_02; 40_40_05 | Firmicutes | Bacilli | Bacillales | Paenibacillaceae |
| Collimonas sp. | 04_04_04; 21_21_04; 21_RT_01 | Proteobacteria | Betaproteobacteria | Burkholderiales | Oxalobacteraceae |
| Dyella japonica | 30_RT_04 | Proteobacteria | Gammaproteobacteria | Xanthomonadales | Rhodanobacteraceae |
| Dyella marensis | 21_RT_04 | Proteobacteria | Gammaproteobacteria | Xanthomonadales | Rhodanobacteraceae |
| Labrys methylaminiphilus | 30_30_08 | Proteobacteria | Alphaproteobacteria | Rhizobiales | Xanthobacteraceae |
| Nocardia coeliaca | 21_RT_05 | Actinobacteria | Actinobacteria | Corynebacteriales | Nocardiaceae |
| Paenibacillus sp. | 21_RT_03 | Firmicutes | Bacilli | Bacillales | Paenibacillaceae |
| Pseudomonas helmanticensis | 04_04_02; 04_04_06; 04_RT_02; 04_RT_05 | Proteobacteria | Gammaproteobacteria | Pseudomonadales | Pseudomonadaceae |
| Pseudomonas protegens | 10_RT_01; 10_RT_02 | Proteobacteria | Gammaproteobacteria | Pseudomonadales | Pseudomonadaceae |
| Pseudomonas rhodesiae-like | 21_21_02; 21_21_06 | Proteobacteria | Gammaproteobacteria | Pseudomonadales | Pseudomonadaceae |
| Rummeliibacillus pycnus | 40_40_04 | Firmicutes | Bacilli | Bacillales | Planococcaceae |
| Rummeliibacillus stabekisii | 50_50_01 | Firmicutes | Bacilli | Bacillales | Planococcaceae |
| Streptomyces lactacystinicus | 30_30_01; 30_30_06; 30_30_07 | Actinobacteria | Actinobacteria | Streptomycetales | Streptomycetaceae |
| Streptomyces mirabilis | 30_30_03 | Actinobacteria | Actinobacteria | Streptomycetales | Streptomycetaceae |
| Variovorax boronicumulans | 30_RT_01; 30_RT_02 | Proteobacteria | Betaproteobacteria | Burkholderiales | Comamonadaceae |
| Variovorax soli | 30_RT_05 | Proteobacteria | Betaproteobacteria | Burkholderiales | Comamonadaceae |
| Viridibacillus arenosi | 40_RT_03; 40_RT_04 | Firmicutes | Bacilli | Bacillales | Planococcaceae |
| Viridibacillus sp. | 40_40_06 | Firmicutes | Bacilli | Bacillales | Planococcaceae |

### 2 Phylum differences seen in alternative datasets

To ask whether the higher growth rates and lower respiration rates of *Firmicutes* comparative to *Proteobacteria* was a phenomenon constrained to our small dataset, or whether it was a more general trend observed between the two phyla, we compared this to the data compiled in two meta-analyses - DeLong et al. [2010] and Smith et al. [2019]. DeLong et al. [2010] compiled data on both active (growth phase) and passive (stationary phase) metabolic rates across a range of bacteria (mainly from Makarieva et al. [2005]), which were corrected to 20°C using an activation energy of 0.61eV. In this dataset, *Proteobacteria* have higher active and passive metabolic rates than *Firmicutes* (active rates Wilcoxon rank-sum test $p = 0.0017$; passive rates Wilcoxon rank-sum test $p = 0.0098$, Fig. S1A), consistent with the work presented in our main text. The DeLong *et. al.* dataset also contains maximum growth rate data (also corrected to 20°C using an activation energy of 0.61eV). However, there is no significant difference between the growth rates of the two phyla in these data (Wilcoxon rank-sum test p-value = 0.66, Fig. S1B). Additionally, we investigated differences in the growth rates of data compiled in Smith et al. [2019]. Given the findings in Smith et al. [2019] of hotter-is-better for mesophiles but not thermophiles, and the small number of thermophilic *Proteobacteria*, these data were restricted to mesophiles ($T_{\text{pk}} < 40.5$°C) and temperature corrected to 20°C based on each strain's individual TPC parameters. In this dataset, there is a significant difference between growth rates, with *Firmicutes* on average higher than *Proteobacteria* (Wilcoxon rank-sum test $p = 0.00035$, Fig. S1C). We also compared the distribution of $T_{\text{opt}}$ for both phyla in the data of Smith et al. [2019] and find that *Proteobacteria* account for much more of the low-temperature strains, whilst *Firmicutes* are more associated with high temperatures (Fig. S1D), this is consistent with our temperature isolation findings (main text Fig. 2A).

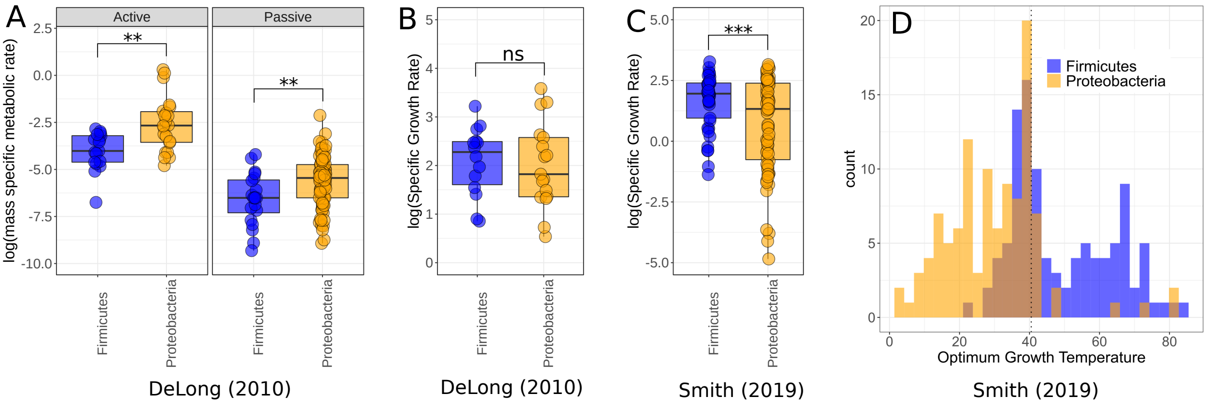


Figure S1: **Comparison of *Firmicutes* and *Proteobacteria* in meta-analysis datasets.**

**A.** Dataset used by DeLong et al. [2010] shows significantly higher active and passive metabolic rates for *Proteobacteria* than *Firmicutes*. Significance determined by Wilcoxon rank-sum tests – ns $p \geqslant 0.05$; * $p < 0.05$; ** $p < 0.01$; *** $p < 0.001$; **** $p < 0.0001$. **B.** The growth rate data used by DeLong et al. [2010] shows no significant difference between the phyla. **C.** The growth rate data from Smith et al. [2019] does show significantly increased growth rates for *Firmicutes* over *Proteobacteria* however. **D.** Distribution of *Firmicutes* and *Proteobacteria* $T_{\text{opt}}$ from Smith et al. [2019]. *Proteobacteria* account for a large proportion of the low temperature strains, whilst *Firmicutes* dominate the high temperatures. Dotted line marks 40.5°C, a cut-off between mesophiles and thermophiles.

### 3 Phylogenetic reconstruction

We used 16S sequences to build a phylogeny in order to investigate the evolution of thermal performance across the isolated bacterial taxa. Sequences were aligned in MAFFT (v7.205) using the default settings. From this alignment 100 trees were inferred in RAxML (v8.1.1), using a GTR-gamma nucleotide substitution model. The tree with the highest log-likelihood was taken and time-calibrated using PLL-DPPDiv, which estimates divergence times using a Dirichlet Process Prior [Heath et al., 2012]. DPPDiv requires a rooted phylogeny with the nodes in the correct order, however RAxML by default produces an unrooted tree. Therefore, we included an archaeal sequence in our 16S alignment (*Methanospirillum hungatei*, RefSeq accession NR_074177) and used this as an outgroup in our RAxML run. This gives a tree rooted at the outgroup, which we checked for correct topology using TimeTree [Kumar et al., 2017] as a reference. We derived calibration nodes from TimeTree [Kumar et al., 2017] (see supplementary table S2) and performed two DPPDiv runs for 1 million generations each, sampling from the posterior distribution every 100 generations. We ensured that the two runs had converged by verifying that each parameter had an effective sample size above 200 and a potential scale reduction factor below 1.1. We summarised the output of DPPDiv into a single tree using the TreeAnnotator program implemented in BEAST [Bouckaert et al., 2019]. We then dropped the outgroup tip to give a time-calibrated phylogeny of our bacterial 16S sequences only, which was used for further analysis.

### 4 Alternative model of trait evolution

In the main text we test phylogenetic heritability of $T_{\text{opt}}$ using Pagel's $\lambda$ metric [Pagel, 1999]. Blomberg's $K$ is another metric which is also widely used to infer phylogenetic heritability [Blomberg et al., 2003, Münkemüller et al., 2012]. Blomberg's $K$ calculates the phylogenetic signal strength as the ratio of the mean squared error of the tip data and the mean squared error of the variance-covariance matrix of the given phylogeny, under the assumption of Brownian motion (BM) [Münkemüller et al., 2012]. $K = 1$ indicates taxa resembling each other as closely as would be expected under a BM model, $K < 1$ indicates less phylogenetic signal than expected under BM and $K > 1$ indicates more phylogenetic signal than expected and thus a substantial degree of trait conservatism [Blomberg et al., 2003]. Under a Brownian motion model of trait evolution, Pagel's $\lambda$ is expected to perform better than $K$, which may itself be better utilised for simulation studies [Münkemüller et al., 2012]. Previous work suggests that $T_{\text{pk}}$ is likely to evolve in a Brownian motion manner in prokaryotes [Kontopoulos et al., 2020], making $\lambda$ a more appropriate metric for these data than $K$. Furthermore, $\lambda$ is potentially more robust to incompletely resolved phylogenies and is therefore likely to provide a better measure than $K$ for ecological data in incomplete phylogenies [Molina-Venegas and Rodríguez, 2017]. Therefore, we use $\lambda$ in the main text as likely the more appropriate metric for our data. Here, for the sake of completeness, we also test for phylogenetic heritability using $K$. We find that Blomberg's $K$ metric also produces a strong signal of phylogenetic heritability in our $T_{\text{opt}}$ data ($K = 0.979$, $p < 0.001$). This is qualitatively the same result as the $\lambda$ test given in the main text.

### 5 Timetree calibration nodes

As part of the time-calibration of our RAxML tree, we constrained the dates of particular nodes in our tree, based on the divergence times of different clades as estimated by TimeTree [Kumar et al., 2017]. Details of calibration nodes used are given in table S2.

Table S2: **Details of time-tree calibration nodes.** We constrained the time-calibration of our RAxML tree based on estimated divergence times from TimeTree [Kumar et al., 2017].

| Taxa A | Taxa B | Min divergence time (MYA) | Max divergence time (MYA) |
|---|---|---|---|
| Bacteria | Archaea | 4,290 | - |
| Pseudomonas | Bacillus | 3,100 | 3,254 |
| Pseudomonas | Labrys | 1,053 | 3,135 |
| Pseudomonas | Collimonas | 1,053 | 3,135 |
| Collimonas | Variovorax | 1,271 | - |
| Arthrobacter | Streptomyces | 1,420 | 1,870 |
| Arthrobacter | Bacillus | 3,100 | 3,254 |
| Bacillus | Brevibacillus | 1,734 | 2,398 |
